# Spontaneous suppressors against debilitating transmembrane mutants of *Ca*Mdr1 disclose novel interdomain communication via Signature motifs of the Major Facilitator Superfamily

**DOI:** 10.1101/2022.03.23.485484

**Authors:** Suman Sharma, Atanu Banerjee, Alexis Moreno, Archana Kumari Redhu, Pierre Falson, Rajendra Prasad

## Abstract

The Major Facilitator Superfamily (MFS) includes multiple families of proteins operating as uniporters, symporters and antiporters for a wide spectrum of substrates. Among them, the multidrug resistance-1 drug:H^+^ antiporter *Ca*Mdr1 from *Candida albicans* is responsible for the efflux of structurally-diverse antifungals. MFS share a common fold of 12-14 transmembrane helices (TMHs) forming two N- and C-domains. Each domain is arranged in a pseudo symmetric fold of two tandems of 3-TMHs that alternatively expose the drug-binding site towards the inside or the outside of the yeast to promote drug binding and release. MFS show a high primary structure diversity and few conserved Signature motifs, each thought to have a common function in the superfamily, although not yet clearly established. Here, we provide new information on these motifs by having screened a library of 64 drug transport-deficient mutants and their corresponding suppressors spontaneously rescuing the deficiency. We found that five strains recovered the drug-resistance capacity by expressing *Ca*Mdr1 with a secondary mutation. The pairs of debilitating/rescuing residues are distributed either in the same TMH (T127A_TMH1_->G140D_TMH1_) or 3-TMHs repeat (F216A_TMH4_->G260A_TMH5_), at the hinge of 3-TMHs repeats tandems (R184A_TMH3_->D235H_TMH4_, L480A_TMH10_->A435T_TMH9_), and finally between the N- and C-domains (G230A_TMH4_->P528H_TMH12_). Remarkably, most of these mutants belongs to the different Signature motifs, highlighting a mechanistic role and interplay thought to be conserved among MFS. Results point also to the specific role of TMH11 in the interplay between the N- and C-domains in the inward- to outward-open conformational transition.

## INTRODUCTION

*C. albicans* is a commensal pathogen that can lead to serious infections, particularly under compromised immunity in human host. Amongst the various strategies adopted by the yeast to resist the antifungal onslaught, elevated drug efflux contributes significantly to an expeditious advent of antifungal resistance (White *et al*. 2002). This reduced intracellular accumulation of drugs in *Candida* is predominantly accredited to *Ca*Cdr1 and *Ca*Cdr2 belonging to the ATP-binding cassette (ABC) transporter proteins and the MFS protein *Ca*Mdr1 (Prasad *et al*. 1995; Sanglard *et al*. 1995; White 1997; Lopez-Ribot *et al*. 1998; Harry *et al*. 2002).

MFS is extensively distributed in many domains of life (Saier Jr *et al*. 2014; Finn *et al*. 2016), forming the broadest and most renowned superfamily of secondary transporters that so far gathers 105 families (http://tcdb.org/superfamily.php; (Wang *et al*. 2020)). MFS members operate as uniporters, symporters and antiporters. They are unique in exhibiting a wide spectrum of substrates (Lee *et al*. 2016). Symporters and antiporters take the advantage of the electrochemical potential of co-transported solute or ion whereas uniporters mediate the facilitated diffusion of a single type of substrate along their concentration gradient (Shi 2013). Most of the MFS proteins share a common scaffold for all members of the family, made of 12-14 TMHs (Law *et al*. 2008). The genome of *C. albicans* features 95 MFS proteins divided into 17 families (Gaur *et al*. 2008), among them the Drug:H^+^ Antiporter 1 (DHA1) family which contains 22 transporters including *Ca*Mdr1utilizing an electrochemical gradient of protons to facilitate the transport of cargo against its concentration gradient across the membrane (Ben-Yaacov *et al*. 1994). Among the *C. albicans* MFS proteins only *Ca*Mdr1 has been linked to a resistance phenotype towards azoles antifungals, as well as several unrelated drugs like 4-nitroquinoline–N-oxide, cycloheximide, benomyl, methotrexate and cerulenin (Ben-Yaacov *et al*. 1994; Gupta *et al*. 1998; Kohli *et al*. 2001; Prasad and Kapoor 2005).

Majority of structural information have come from prokaryotic MFS transporters and, to a lower extent, from eukaryotic homologs (Huang *et al*. 2003; Yin *et al*. 2006; Thorens and Mueckler 2009; Sun *et al*. 2012; Zheng *et al*. 2013; Yan *et al*. 2013; Jiang *et al*. 2013; Deng *et al*. 2014; Yan 2015; Drew *et al*. 2021a). Twelve-TMHs MFS are made of two N- and C- 6-TMHs subdomains that are organized as two inverted pairs of 3-TMHs bundles (Radestock and Forrest 2011; Madej and Kaback 2013; Yan 2015). MFS display a poor primary structure identity but share few conserved Signature motifs thought to play similar and key roles (Paulsen *et al*. 1996; Monique *et al*. 2000). Motif A (*GxLaD^180^rxGrkx_3_I,* referring to the *Ca*Mdr1 sequence numbering) is located in the cytoplasmic loop between TMH2 and TMH3. It is supposed to be involved in the inward/outward conformational change (Monique *et al*. 2000) and later found to be stabilizing the outward-facing conformation of YajR (Jiang *et al*. 2013) by salt-bridges either in the A motif or with adjacent regions (Kakarla *et al*. 2017; Kumar *et al*. 2020a). Motif B (*Ix_3_R^215^x_2_qGxga_2_*) is located in the external leaflet of TMH4. It contains an arginine residue inferred to be involved in proton transfer which indeed we confirmed for *Ca*Mdr1 (Redhu *et al*. 2016). Motif C (*gx_3_G^260^Px_2_G_2_xI*) is positioned in the external leaflet of TMH5. It displays two Gx_3_G motifs, known to stabilize helix-helix association in membrane proteins through interaction with bulky side-chain residues (Russ and Engelman 2000). Mutation of these glycine residues in TetA (Ginn SL *et al*. 2000) and *Ca*Mdr1 (Ritu *et al*. 2007) was indeed critical. Motif D (*lgx_5_P^139^vxP*) in TMH1 and motif G (*Gx_3_GPL^512^*) in TMH11 are exclusive to the 12-(motif D) and 12-14-TMH (motif G) families, respectively (Paulsen *et al*. 1996). Both motifs have been so far poorly investigated, but alanine scanning of the corresponding regions of *Ca*Mdr1 (*MGSAVYTP^139^GIE* and *IASVFPL^512^*) showed that residues M132, Y137, V506, A508, P512 and L513 are indeed structurally or functionally critical (Redhu *et al*. 2018).

The *E. coli* lactose/H^+^ symporter LacY has been the most extensively studied among the MFS. Its X-ray structures in inward-open and ligand-bound occluded conformations provided a prototype for understanding the transport mechanism (Shuman 1981; Abramson *et al*. 2003; Guan *et al*. 2007; Kumar *et al*. 2018). Several elegant structural studies in the last decade ensured the visualization of multiple substrate-bound transporter conformations from which a general alternative access mechanism of transport has been deduced. Mechanistically, each protein has a single substrate-binding cavity in the center of the membrane domain. The switch between the inward-open and outward-open conformations at the N- and C-domains interface exposes this cavity to either side of the membrane. N- and C- moieties contribute asymmetrically to form the substrate-binding pocket in symporters and facilitators whereas they contribute equally in antiporters (Yan 2015; Drew *et al*. 2021b). The presence of aromatic residues in the cavity prevents the exposure of the substrate to the cytosolic or extracellular sides (Yan 2015). Detailed biochemical, biophysical, and structural investigations of the MFS antiporters MdfA, EmrD, YajR and SotB from *E. coli* and LmrP from *L. lactis* revealed that the substrate - H^+^ coupling mechanism involves the sequential binding and release of substrate and proton. Both halves of the protein move correlatively similar to a rocker switch, arbitrated by salt bridge formation and breakage during the transport cycle (Jiang *et al*. 2013; Wisedchaisri *et al*. 2014; Zhang *et al*. 2015; Tan and Wang 2016; Debruycker *et al*. 2020; Xiao *et al*. 2021; Drew *et al*. 2021b). Further studies suggest that although proton translocation and substrate transport occur in distinct sites, they always compete for protein binding. Consequently, protonation leads to conformation changes of the protein that facilitate substrate uptake from intracellular side (inward-open conformation) whereas deprotonation destabilizes the substrate-bound state of the protein and eventually leads to substrate release on the extracellular side (outward open conformation) (Yin *et al*. 2006; Schaedler and Van Veen 2010; Fluman *et al*. 2012; Heng *et al*. 2015; Wu *et al*. 2020; Drew *et al*. 2021b).

Our groups have been focusing on the functional aspects of *Ca*Mdr1, mainly by subjecting it to site-directed mutagenesis and homology modeling (Ritu *et al*. 2007; Kapoor *et al*. 2009; Mandal *et al*. 2012; Redhu *et al*. 2016, 2018). These studies firstly highlighted the role of a central cytoplasmic loop (CCL or ICL_3_) in establishing contact between the protein and the plasma membrane (Mandal *et al*. 2012). Then, site-directed mutagenesis guided by prediction of critical residues based on information theoretic measures allowed us to identify several functionally relevant residues (Kapoor *et al*. 2010). Finally, the systematic replacement by alanine of the 252 residues forming the membrane domain of *Ca*Mdr1 revealed 84 residues critical for drug efflux that we categorize depending of their type and impact on either, *Expression* (addressing issues), *Structure* (typically glycine, proline, alanine residues), interaction with *Lipids* (*i.e.* facing the membrane), *Mechanism* (buried residues but not facing the drug-binding cavity), substrate *Binding* (buried residues facing the drug-binding pocket), and *Polyspecificity* (same as B but displaying substrate selectivity). Notably, the spatial organization of residues belonging to the two last groups draw the structural features of the drug polyspecificity characterizing such proteins (Redhu *et al*. 2018).

These studies provide a fair understanding of drug-protein interaction but lack information about the dynamics of the mechanism and interaction between critical residues. To explore these aspects, we took advantage of our *in-house* library of critical mutants that we subjected to the suppressor genetics strategy. This led to 16 strains recovering a resistance to antifungals from initial transport-sensitive mutants belonging to the *Structure/Lipids*, *Mechanism*, *Binding* and *Polyspecificity* groups. Among them, only strains expressing mutants from the two first groups led to stable and intragenic secondary mutations which, strikingly, target the conserved MFS motifs, delivering new information on short and long ranges dynamics of the antiporter and role of these motifs.

## RESULTS & DISCUSSION

### Generation of *Ca*Mdr1 drug-sensitive suppressors

Using our 84 critical *Ca*Mdr1 alanine-mutants library (Redhu *et al*. 2018), we forced yeast expressing 64 of them (see **Supplementary Tables 1-2** in the Methods section) to grow on media containing toxic concentrations of either cycloheximide (CHX), 4-nitroquinoline (4-NQO) or fluconazole (FLC) (**Sup. Fig. 1**). Several rounds of screens selected newly drug-resistant colonies from 16 different strains (**Table 1**), corresponding to initial mutants that originated from most of the categories previously defined in respect of their initial impact on the antiporter (Redhu *et al*. 2018), *Lipids* for F216, Y408 and I448, *Structure* for G230, P257 and L480, *Mechanism* for I123, T127 and R184, *Binding*/*Polyspecificity* for W249, Y365, Y369, F371, F474, Q478 and V506 (**Sup. Fig. 1**). When assessing the restored drug-resistance phenotype of these strains both by the microdilution method and MIC_80_ determination, most of the yeast expressing mutants belonging to the *Binding* group, W249A, Y365A, Y369A, F371A and Y408A, together with I123A, did not sustain their growth in the presence of drugs, indicating a transient effect gradually lost (**Table 1, Sup. Fig. 1**). Sequencing *CaMDR1* in the 10 other strains showed that those expressing P257A, I448A, F474A, Q478A and V506A mutants did not have a secondary mutation inside the gene, implying an intergenic phenotypic effect (**Table 1**). Interestingly, all these residues are positioned at the interface between the inner and outer leaflets of the membrane (**Sup Fig. 1**). Finally, five strains displayed a secondary mutation along with the primary mutation within the *Camdr1* gene, T127A-G140D, R184A-D235H, F216A-G260A, G230A-P528H and L480A-A435T. All these primary alanine mutants are restricted to the *Structure*/*Lipids* (F216, G230, L480) and *Mechanism* (T127, R184) groups.

**Table 1.** Position, location and drug profiling of alanine mutants and suppressors. Phenotypes of alanine mutants and suppressor colonies are represented as TS (Total Susceptibility), SS (Selective Susceptibility), TR (Total Resistance), SR (Selective Resistance), PR (Partial Resistance).

|  | Primary mutant |  |  |  | Suppressor |  |
| --- | --- | --- | --- | --- | --- | --- |
|  | mutation | location | phenotype | Function<br>(Redhu et al. 2018) | Drug<br>leading to<br>resistance | phenotype |
| Transiently resistant strains | I123A | TMH-1 | SS | drug/H<sup>+</sup> antiport mechanism | CHX,<br>4-NQO | TS |
|  | W249A | TMH-5 | TS | drug binding and transport | CHX | TS |
|  | Y365A | TMH-7 | TS | drug binding and transport | CHX,<br>4-NQO | TS |
|  | Y369A | TMH-7 | TS | drug binding and transport | CHX | PR |
|  | F371A | TMH-7 | TS | drug binding and transport | CHX,<br>4-NQO | PR |
|  | Y408A | TMH-8 | TS | exposed to lipid interface | 4-NQO | PR |
| Resistance phenotype<br>(intergenic mutation) | P257A | TMH-5 | TS | structural integrity | CHX | TR |
|  | I448A | TMH-9 | TS | Crucial for plasma membrane localization | CHX,<br>4-NQO | TR |
|  | F474A | TMH-10 | TS | drug binding and transport | CHX | SR |
|  | Q478A | TMH-10 | TS | drug binding and transport | CHX | SR |
|  | V506A | TMH-11 | TS | Crucial role in polyspecific substrate binding and transport | CHX | TR |
| Resistance phenotype<br>(intragenic mutation) | T127A | TMH-1 | TS | drug/H<sup>+</sup> antiport mechanism | CHX | SR |
|  | R184A | TMH-3 | TS | drug/H<sup>+</sup> antiport mechanism | CHX | TR |
|  | F216A | TMH-4 | TS | exposed to lipid interface | CHX | TR |
|  | G230A | TMH-4 | TS | structural integrity | CHX | SR |
|  | L480A | TMH-10 | TS | structural integrity | 4-NQO | TR |

**Figure 1:**
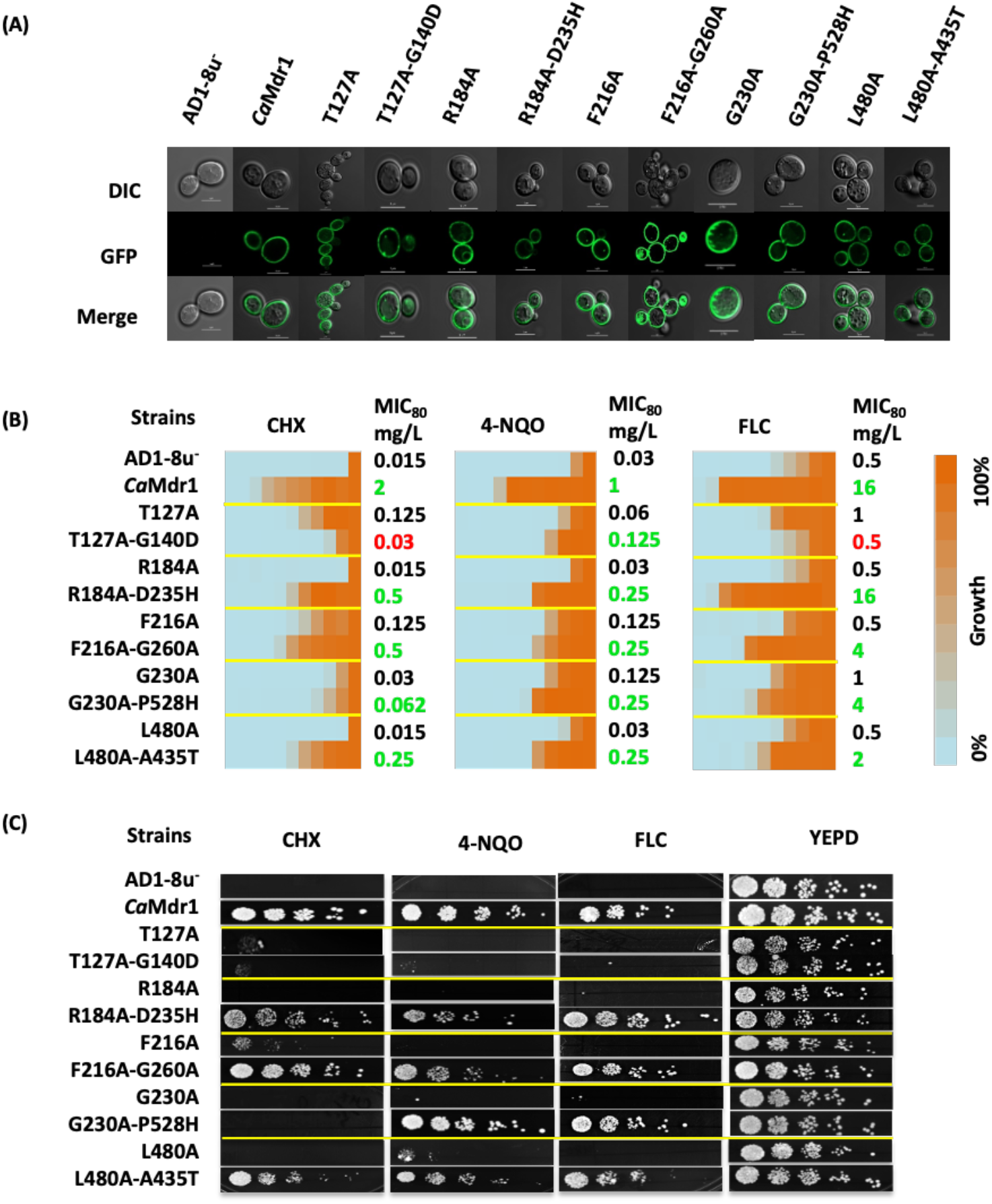
Cell localization and drug resistance profile of primary alanine mutants and their corresponding suppressor mutants. (**A**) *Ca*Mdr1 suppressor mutants localized by confocal microscopy. (**B**) Drug heat map and MIC_80_ values for the corresponding strains. A 2-fold dilution was applied to 8 mg/L CHX and 4-NQO and 32 mg/L FLC. (**C**) Five-fold serial dilution spot assays of the same strains done on solid YEPD medium added of either 0.15 mg/L CHX, 0.15 mg/L 4- NQO or 0.8 mg/L FLC. Data have been collected after 48-h incubation at 30 °C from 3 independent experiments.

We re-constructed these 5 pairs of mutants in the WT *CaMDR1-*GFP gene to exclude extragenic effects and overexpressed them in the *S. cerevisiae* AD1-8u^−^, a host strain that has proven to be an excellent heterologous system for drug transporter overexpression (Decottignies *et al*. 1998; Erwin *et al*. 2007). We designated these strains as Mdr1[L480A-A435T]-GFP, Mdr1[R184A-D235H]-GFP, Mdr1[F216A-G260A]-GFP, Mdr1[T127A-G140D]-GFP and Mdr1[G230A-P528H]-GFP. Using confocal microscopy, GFP fluorescence confirmed for the proper localization at the plasma membrane of each protein (**Fig. 1A**).

Those strains were then subjected to drug susceptibility tests towards CHX, 4-NQO and FLC (**Fig. 1B-C**). MIC_80_ values and spot assays showed that suppressors strains expressing the L480A-A435T, R184A-D235H and F216A-G260A *Ca*Mdr1 variants grow in the presence of drugs with up to 30-fold MIC_80_ increase as compared to their respective primary alanine mutants. Although initially isolated through resistance to CHX, the strain expressing the T127A-G140D variant hardly maintained such resistance, together with that towards FLC. However, it remained slightly more resistant to 4-NQO than the parental strain. Finally, the yeast expressing the G230A-P528H variant displayed the same sensitivity pattern to CHX than the later but remains 2-4-fold more resistant to 4-NQO and FLC.

### Most of the drug-sensitive mutants and suppressors locate within the conserved Signature motifs of DHA1 MFS

As suggested by our 3D model, primary and secondary mutants distribute along TMH1, 3, 4, 5, 9,10 and 12 (**Fig. 2**), either belonging to the same TMH (1) for T127A and G140D, or most often in two different TMHs. In the inward-facing 3D model of *Ca*Mdr1, those pairs are either close for R184A_TMH3_-D235H_TMH4_, F216A_TMH4_-G260A_TMH5_ and L480A_TMH10_-A435T_TMH9_, or far from each other for G230A_TMH4_-P528H_TMH12_ (**Fig. 2**). Seven over the ten modified residues are in the N-domain of *Ca*Mdr1.

**Figure 2.**
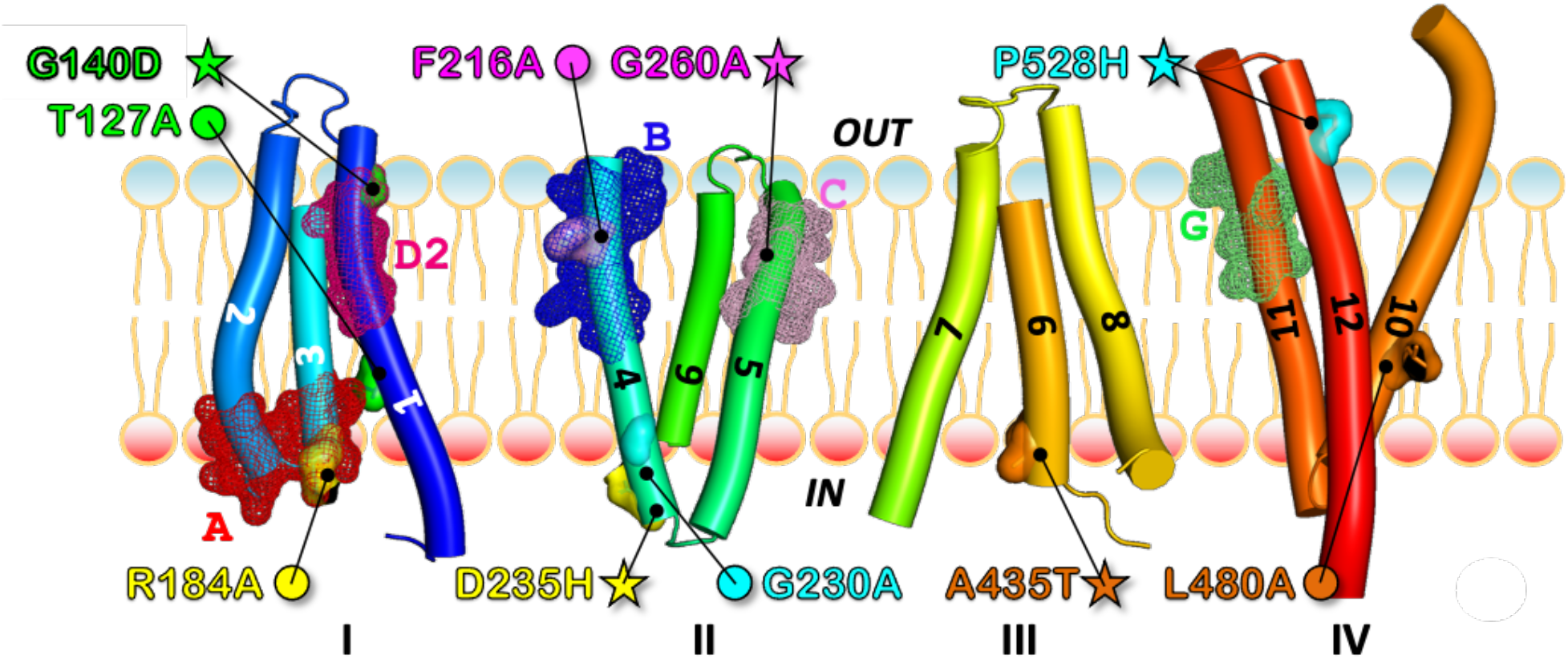
Localization of the couples of primary-debilitating and secondary-rescuing transport mutants of *Ca*Mdr1 in respect of MFS Signature motifs and internal structural repeats. 3D model of *Ca*Mdr1 in inward facing conformation displayed in respect of the four structural repeats (Radestock and Forrest 2011) and the conserved A, B, C, D2 and G signature motifs of the proton-dependent multidrug efflux systems (Paulsen *et al*. 1996). See supplementary figures 2 and 3 for details. Primary-debilitating (circles) and secondary-rescuing (stars) mutants are shown in surface and sticks and indicated with the same color for each couple. Conserved motifs are shown in mesh.

Remarkably, looking at these mutants in respect of the DHA1 MFS subfamily motifs (Paulsen *et al*. 1996) revealed that G140 belongs to motif D2, R184 to motif A, F216 to motif B, G260 to motif C, and that T127 and P528 are close to motifs D2 and G, respectively (**Fig. 2, Sup. Fig. 2-3**). As introduced above, conserved residues of these motifs are thought to have a common key mechanistic role within MFS, which is in line with the fact that the present restoration process was finally successful for residues of the *Mechanism* and *Structure* groups (**Table 1, Sup. Fig. 1**).

### Localization of sensitive and suppressor mutants in the outward-facing conformation of *Ca*Mdr1

To get a better view of the positional significance of mutant and suppressor couples we generated the 3D model of the outward-facing conformation of *Ca*Mdr1, covering residues 110 to 544. We built it on YajR, an *E. coli* proton-driven MFS antiporter crystallized in this conformation (Jiang *et al*. 2013), PDB code 3WDO. Primary sequences alignment of *Ca*Mdr1 and YajR (UniprotKB Q9URI1 and P7726, respectively) using AlignMe (Stamm *et al*. 2014) showed 10% sequence identity and 63% matched position. The alignment was used to manually superimpose the inward-facing model of *Ca*Mdr1 with the crystal structure of YajR with Pymol (Version 2.5.0 Schrödinger, LLC.). The alignment was then submitted to Modeller (Webb and Sali 2021) that generated 20 models among which the more representative was selected manually. The final outward-facing model (**Fig. 3A, right panel**) displays a reorientation towards the external side of the membrane of the N- and C-moieties, exposing the drug-binding pocket to the extracellular space. Comparison with the inward-facing model (**left panel**) shows the spatial distribution of conserved Signature motifs. Most of them are clustered in (A, B, C, D2) or close to (G2) the N-ter moiety. The models also highlight the remarkable alignment of B and C motifs with D2 in between. We also took advantage of the recent developments of 3D modelling offered by AlphaFold (Jumper *et al*. 2021) to compare this model with ours, and we found it close to our inward-open model (**Sup. Fig. 4**).

**Figure 3.**
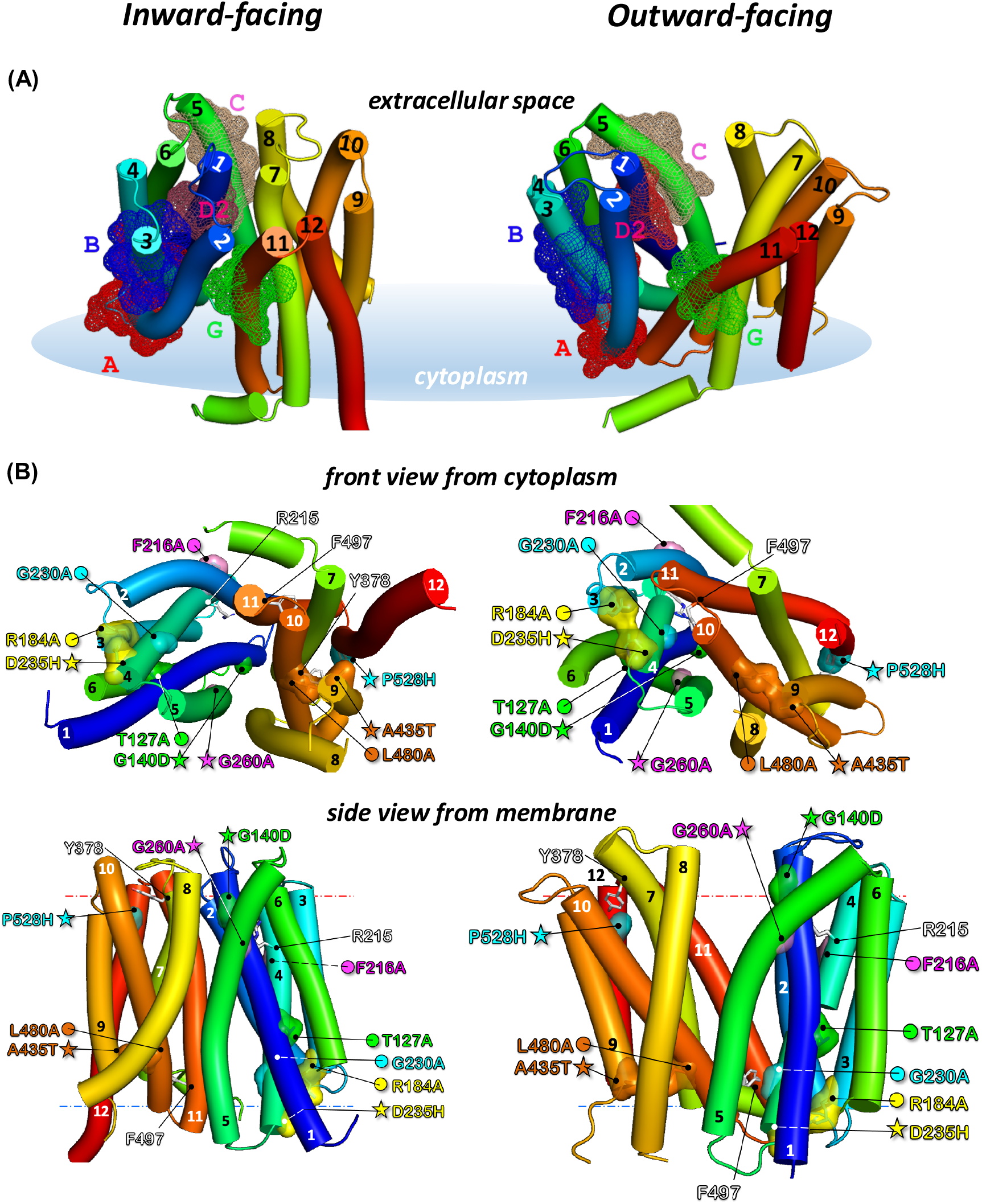
Position of conserved MFS Signature motifs and location of primary mutant and secondary suppressor couples in the inward- and outward-facing models of *Ca*Mdr1. (**A**) Left, inward-facing conformation based on GlpT sequence (Redhu *et al*. 2016), optimized with Modeller. Right, outward-facing conformation based on YajR crystal structure (Jiang *et al*. 2013). Models are displayed in solid cartoon (Pymol v2.5.0), colored from the N-blue to the C-red ends. Signature motifs are defined in Fig. 2. (**B**) Front and side views of the inward- and outward-facing models with position of primary debilitating (circles) and secondary restoring (stars) mutants. Residues are colored by couples. Residues R215, Y378 and F497 are discussed in the text. Blue and red dotted lines indicate cytoplasmic and extracellular membrane limits as defined by the PPM server (https://opm.phar.umich.edu/ppm_server).

Checking the distribution of the 84 critical residues in respect of their category in this new conformation (**Sup. Fig. 5**) shown that residues belonging to the *Binding* and *Polyspecificity* groups (green and orange, respectively) remain facing the drug-binding cavity (right panels) and those interacting with lipids (pink) are still facing them, while those involved in the mechanism (blue) remain mainly clustered in the N-moiety. In addition, residues conferring polyspecificity (orange) are still localized at the periphery of those involved in substrate binding (green), which strengthens our previous finding that such pattern is a molecular feature of MDR pumps polyspecificity (Redhu *et al*. 2018; Banerjee *et al*. 2021).

With these models, we positioned the different primary sensitive and secondary resistant mutant and suppressor couples (**Fig. 3B**). Comparing their location in our inward-open model and the AlphaFold one, we observed that the mutants and revertants are in the same place or close in both models. One main difference is the larger loop linking TMH11 and 12 predicted by AlphaFold, displacing P528 in that loop at the top of the protein. The residue remains however well oriented towards Y378 (described below) and close to it (**Sup. Fig. 4**).

### Local compensations restore 3-TMHs repeats tandems interactions in N- and C-domains and highlight the role of motif A

Looking first to mutant and suppressor couples spatially close, namely R184A-D235H and L480A-A435T in the N- and C-domains, respectively (**Fig. 3B**), revealed a series of common features, *i*) each couple is perfectly symmetrically positioned in the axis of the membrane in their respective domain, *ii*) they are close to the cytoplasm side of the membrane domain, *iii*) each residue belongs to a specific 3-TMHs repeat, I for R184, II for D235 and III for A435 and IV for L480 (**Fig. 2**), and *iv*) they remain close whatever the inward-facing or outward-facing conformation.

R184 brings a positive charge to the motif A where it belongs (**Figs. 2, 3 and Sup. Fig. 3**) and which is engaged in a salt bridge with D235, both in inward- and outward-facing models of *Ca*Mdr1 (**Fig. 3, Sup. Fig. 6A**). The salt bridge contributes to stabilize the interaction between TMH3 and TMH4 and consequently between 3-TMHs repeats I and II (**Fig. 2**). D235 is rather well conserved (**Sup. Fig. 3**) and indeed critical as the D235H single mutant that we generated confers a full sensitivity to drugs to the yeast expressing the corresponding *Ca*Mdr1 variant (**Sup. Fig. 7AB**). This observation strengthens the hypothesis of a stabilizing role of the salt bridge, as also concluded for the corresponding residues of TetA(B), R70 and D120 (Someya *et al*. 2000). However, this does not exclude a specific role of the positive charge provided by R184 since the compensation process privileges the restoration of a positive charge with the D235H substitution in the background of R184A. However, this option does not seem relevant since when exploring the pH sensitivity of those mutants in the presence or absence of 4-NQO did not reveal any significant dependency towards this parameter (**Sup. Fig. 7CD**).

In the C-domain the L480A mutation in TMH10 is compensated by A435T in TMH9 (**Fig. 3**), for which inward- and outward-facing models of *Ca*Mdr1 suggest that both residues interact together within a local network of hydrophobic interactions involving aliphatic (I412_TMH8_, I434_TMH9_, V437_TMH9_, I476_TMH10_, M484_TMH10_) and aromatic (Y408_TMH8_, F477_TMH10_, F481_TMH10_) residues (**Sup. Fig. 6B**). The size reduction of the aliphatic tail of L480 to alanine and the central location of the couple of residues seems therefore enough to weaken such network. This hypothesis is strengthened by the effect of the A435G mutation that we found previously fully deleterious for multi-drug efflux (Redhu *et al*. 2018), contrarily to the single A435T mutation that we generated (**Sup. Fig. 6C**). Introducing the two mutations L480A and A435T in the inward- and outward-facing models of *Ca*Mdr1 shows that T435 may be well positioned to generate a H-bond with the Sulphur atom of M484, one helix turn below L480 on TMH10 (**Sup. Fig. 6B**), which may contribute to restore the interaction lost with the L480A mutation.

Altogether these data suggest that the regions in which the two couples of mutants and suppressors are located grant a stable interaction between their respective TMHs, allowing to synchronize the corresponding pairs of 3-TMHs repeats in each domain along the drug translocation process. Well conserved, the motif A may indeed play this role in MFS proteins. This can be also the case for the regions encompassing A435 and L480, although a lower conservation level compensated by an enrichment in aliphatic and aromatic residues (**Sup. Fig. 3**). The natural abundance and variability of such residues reduce the requirement of identity since hydrophobic properties are preserved, which therefore masks a potential signature.

### Distant compensations in the N-domain highlight motifs A, B and D2 interplay

Looking at the mutant and suppressor couple F216A-G260A reveals that both residues belong to the same 3-TMHs repeat II and that they are part of motifs B in TMH4 and motif C in TMH5, respectively (**Fig. 2**). F216 and G260 stand in the same plane in the extracellular leaflet of the membrane, at a distance of ~20 Å from each other, both in inward- and outward-facing models (**Fig. 3B**). F216 is rather well conserved in motif B (**Sup. Fig. 3**), together with F220 positioned one helix turn below, that we also previously identified critical when mutated into alanine (Redhu *et al*. 2018). Inward and outward-facing models of *Ca*Mdr1 show that both aromatic residues face lipids (**Fig. 3B**), suggesting that their replacement by an alanine may increase the level of freedom of the corresponding segment of TMH4 in the outer leaflet. This may alter a precise location of R215 that follows F216 on the opposite face of TMH6 and points to TMH1 in the center of the membrane domain (**Fig. 3**) where it plays the main role in proton antiport (Redhu *et al*. 2016). In the background of the F216A mutation, drug efflux is restored through the G260A substitution in TMH5, pointing at the same level than R215 to the other side of TMH1 (**Fig. 3**). Member of the motif C, G260 is the central glycine residue of a double glycine motif G^256^-X-X-X-G^260^-X-X-X-G^264^ (**Sup. Fig. 3**), which is typical of membrane helix-helix interaction (Russ and Engelman 2000) and faces the outer leaflet part of TMH1. The glycine motif contributes to the tight and constant interaction between TMH4 and TMH1. Glycine substitution to alanine is possible in such motif, and indeed we previously found that such substitution is not deleterious (Redhu *et al*. 2018). But a methyl group, bulkier than a proton, may probably push the outward leaflet part of TMH1 towards TMH5, which contributes to reconnect R215 to the proton translocation network.

The region of TMH1 to which R215 and G260 point to corresponds to the motif D2 (**Figs. 2, 3**) that we found here targeted by the genetics strategy with G140D restoring the drug resistance lost with the T127A mutation. T127 precedes the motif D2 while G140 is located at its C-ter end (**Sup. Fig. 3**). G140 faces G260 in the models of *Ca*Mdr1 and its substitution by an aspartic residue may produce the same repositioning effect than the G260A mutation but also contribute to reconnect the proton translocation network.

Altogether, these data show that motifs B, C and D2 constitute a bundle finely adjusted to synchronize the motion of TMH1, 4 and 5 for granting proton translocation during drug efflux.

### Long-range compensation between the N- and C-domains

The pair G230A-P528H constitutes a striking event of long-range compensation. G230 is located in C-end of TMH4 within the 3-TMHs repeat II of the N-domain, close to the cytoplasmic side of the protein and oriented towards the drug-translocation pathway. P528 is symmetrically located about 40 Å faraway at the external face of the membrane in N-end of TMH12 in the 3-TMHs repeat IV of the C-domain (**Figs. 2, 3B, 4**). TMH11 seems to be the most direct link between the couple of residues (**Fig. 4**) and, interestingly, carries the G motif in the outer leaflet region of the TMH (**Fig. 2**). Inward-open and outward-open models of *Ca*Mdr1 suggest that TMH11 undergoes a large movement by which its N-Ter comes close to G230 in the outward-open conformation (**Fig. 4A**). Here, residue F497 in TMH11 may be as close as G230 thanks to the empty space provided by the absence of lateral chain of G230, as in a glycine motif. The distance increases up to about 20 Å when coming back to the inward-facing state (**Fig. 4**). This interaction may therefore contribute to the stabilization of the outward-facing state, weakened or hampered by the G230A substitution that locally increases the steric hindrance.

**Figure 4.**
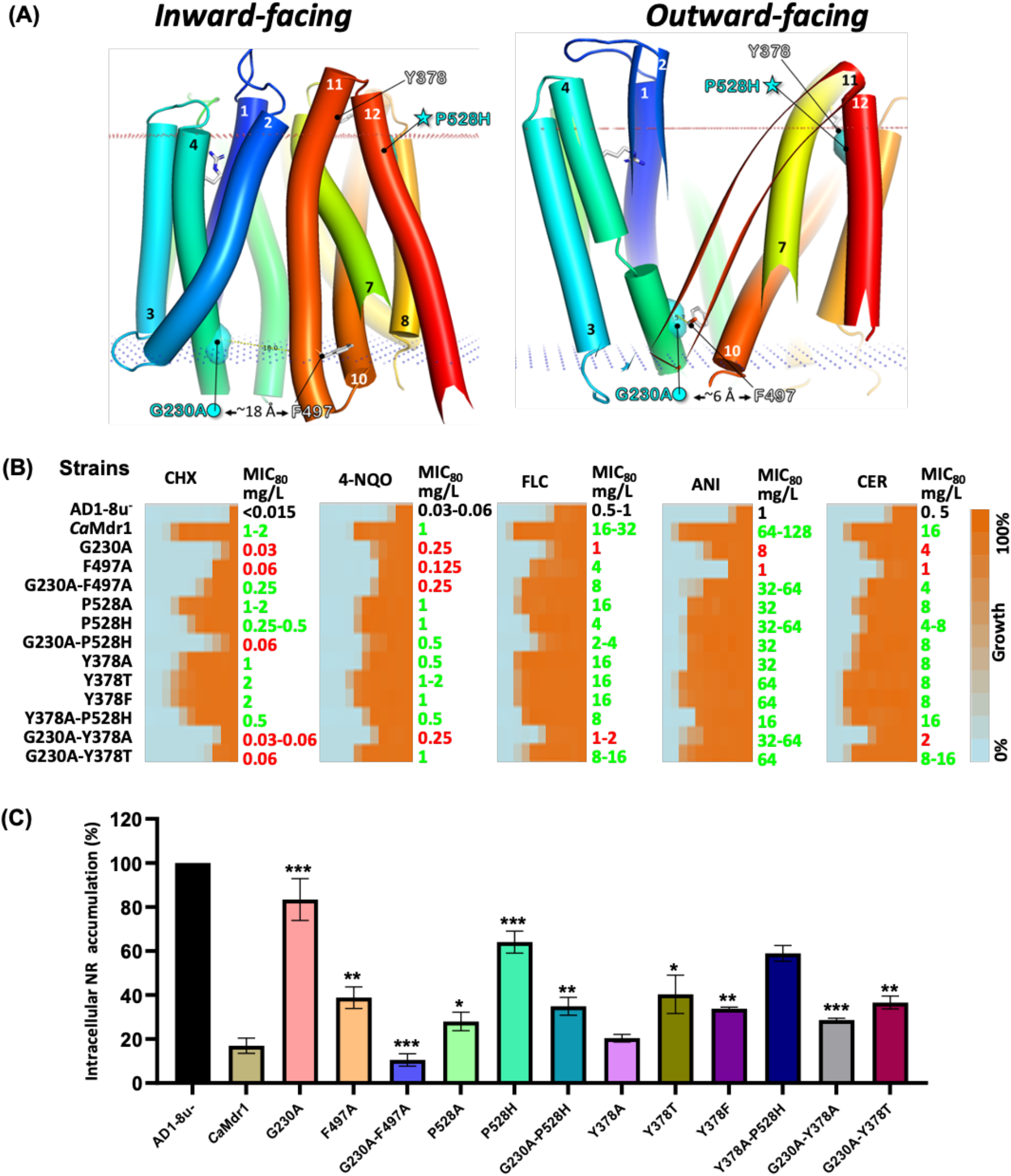
Short- and long-range interactions involving positions 230, 378, 497 and 528. (**A**). Cartoon representation of inward- and outward-facing models of *Ca*Mdr1 with membrane limits. Settings are as in Fig. 3. TMH7, 11 and 12 have been partially masked. (**B**). MIC_80_ values as described in Fig. 1. Anisomycin (ANI, 10 mg/L) and cerulenin (CER, 4 mg/L) have been added to the screen. (**C**). Nile red (NR) accumulation assays in host AD1-8u-, WT *Ca*Mdr1-GFP and variants. NR accumulation in host strain was set to 100%. Results are the mean of three independent cultures.

According to this scenario, reducing the steric hindrance at the position 497 should have a compensatory effect that we evaluated by generating and studying the effect of the F497A mutation, alone and in the background of G230A (**Fig. 4B**, **Sup. Fig. 8**). Both variants were well expressed and addressed in yeast (**Sup. Fig. 8A**). Liquid (**Fig. 4B**) and solid (**Sup. Fig. 8B**) dilution assays showed that the yeast expressing the F497A variant, except for fluconazole, becomes sensitive to cycloheximide and 4-nitroquinoline, and even higher for anisomycin and cerulenin. These results confirm that reducing the steric hindrance at position 497 has the same deleterious impact than increasing it at the position 230. As expected, the strain expressing the *Ca*Mdr1 variant F497A in the background of G230A recovers a significant level of resistance towards most of antifungals (**Fig. 4B**) and full resistance towards Nile red (**Fig. 4C**). These data thus support the functional proximity and steric complementarity of G230 and F497 suggested by the 3D models together with, unexpectedly, a role of these residues and their interaction in drug selectivity.

Exploring the P528 region with the same strategy (**Fig. 4B, Sup. Fig. 8**), added of accumulation assays carried out with Nile red (**Fig. 4C**), showed that the yeast expressing the *Ca*Mdr1 P528H variant remains significantly resistant to all drugs except Nile red, indicating a limited impact of this mutation when present alone. Replacement by an alanine gave the same result. However, introducing the P528H mutation in the background of the deleterious G230A mutation restores a better level of resistance towards the tested drugs and accumulated significantly lower amounts of Nile red, compared to the yeast expressing the single G230A variant.

Inward- and outward-open models of *Ca*Mdr1 suggest that P528 faces TMH7, 8 and 11 (**Fig. 4, Sup. Fig. 9**). A histidine residue at this position may locally increase the steric hindrance but may also bring polarity and charge which probably contributes altogether to reposition TMH12 in respect to the other TMHs for compensating the handicap introduced with G230A. We noticed that in both models P528 is particularly close (~10 Å, considering Cα, as also in the AlphaFold model (**Sup. Fig. 4**)) to Y378 in TMH7 (**Figs. 3, 4**, **Sup. Fig. 9**) which, interestingly, stands in the well conserved short segment P*h*Y*h* (*h* for hydrophobic) (**Sup. Fig. 3**). To gain further insights in the local and distant regions interplay, we generated a series of single and double mutants and analyzed their impact on *Ca*Mdr1 substrates efflux (**Fig. 4**, **Sup. Fig. 8**). Substituting Y378 by an alanine, threonine or phenylalanine did not produce any significant or big effect on substrates efflux, even in the background of P528H. Addition of the secondary P528H mutation in the background of Y378A was more deleterious. However, an alanine or threonine at position 378 in the background of G230A restored a resistance phenotype, confirming that P528 and Y378 are indeed close. Altogether, these data show that TMH11 tunes the relative positions of the N- and C-domains, mainly for allowing the inward- to outward-open conformational change but with consequences on drug selectivity.

## CONCLUDING REMARKS

In this study we describe that a non-directed mutagenesis process applied to a MDR drug:H^+^ antiporter selects primary and secondary mutants mainly located in few conserved stretches of proteins belonging to the DHA1 MFS family, so-called Signature motifs A, B, C, D2 and G. Despite the functional diversity of the starting 64-mutants library, only mutants initially having a deleterious impact either on the mechanism (corresponding to residues with small or bulkier lateral chain pointing inside the protein but not in the drug-binding cavity (Redhu *et al*. 2018)) or the interaction with lipids (residues with lateral chain facing the membrane (Redhu *et al*. 2018)) allow a rescuing secondary mutation to occur. The privileged distribution of these residues among the Signature motifs suggests therefore that these regions may constitute the mechanistic core of DHA1 MFS. Clustered in the outward leaflet, motifs B, C and D2 may provide the power stroke of the efflux while motifs A and G, symmetrically located in the inner and outer leaflet of the membrane may drive the conformational change allowing drug translocation.

## MATERIALS AND METHODS

### MATERIALS

#### Reagents

All chemicals used on a regular basis were acquired from Himedia, Merck, or SRL Pvt. Ltd. Mumbai, India. Drugs used such as cycloheximide (CHX), 4-nitroquinoline 1-oxide (NQO), fluconazole (FLC), anisomycin (ANI), cerulenin (CER), and fluorescent dye nile red (NR) were manufactured by Sigma-Aldrich Co. (St. Louis, MO). Reagents such as ammonium acetate, PEG (Polyethylene Glycol), LiAc (Lithium Acetate), sodium chloride (NaCl), Tris-HCl, EDTA, and dimethyl sulfoxide (DMSO) were also purchased from Sigma-Aldrich Co. (St. Louis, MO). Oligonucleotides were obtained from Sigma Genosys, India and have been listed in **Supplementary Table 1**.

#### Strains and culture conditions

All the yeast strains were grown either in YEPD (Yeast extract, Peptone and Dextrose) broth or on solid YEPD plates with 2% agar with or without drug treatment according the experimental requirement. Dh5α strain of *Escherichia coli* was used to maintain all plasmids and cells were grown in Luria Bertani media with 100 mg/L ampicillin dosage (Amresco, Solon, USA). Both growth media (YEPD and LB media) and Agar were purchased from HiMedia Laboratories, Mumbai, India. To select yeast transformants, Synthetic defined medium without uracil (SD-Ura^−^) plates were used that composed of 0.67% Yeast nitrogen base without amino acids (Difco, Becton, Dickinson and Company, MD, USA), 0.2% Ura^−^ dropout mix (Sigma Genosys, India), and 2% glucose (Merck) along with 2.5% (w/v) agar from Hi Media laboratories, Mumbai, India. All yeast and bacterial strains stock were prepared using 15% glycerol and stored at −80 °C. **Supplementary Table 2** enlisted all the strains used in this study.

### METHODS

#### Generation and sequence analysis of suppressor mutants

Overnight grown cultures of yeast expressing the *Ca*Mdr1-GFP mutant variants were washed with sterile 0.9% saline. The cells were then homogenously mixed with 25 mL molten YEPD agar to accomplish a final OD_600 nm_ of 10^5^ cells/mL at wavelength and were poured into Petri plates. The filter discs were positioned on the plates once the medium had solidified, and the desired toxic concentration of drugs was deposited on the discs using pipette. Afterwards, yeast were allowed to grow under the selective pressure of its drug substrates for 6-7 days at 30°C. The plates were observed regularly and the colonies appeared within the inhibitory zone were picked up and subsequently validated by passage on drug plates. Further, genomic DNA was extracted from the validated colonies by using glass bead method and *CaMDR1* gene was amplified by performing PCR using Phusion polymerase from New England Biolabs and *Camdr1* full gene primers listed in **supplementary Table 1**, manufactured by Sigma Genosys, India. To detect base alterations that resulted in amino acid substitution, PCR amplicons were sequenced using overlapping primers across the whole *Camdr1* ORF. To avoid errors, the sequencing was done at least twice. The resulted sequences were then analyzed by using Align Me software with aligning of sequences with original gene sequence.

#### Site-directed mutagenesis and yeast transformation

To conduct site-directed mutagenesis, Quick-Change kit was used as indicated by the manufacturer (Agilent Technologies, USA). Full plasmid with *CaMDR1* gene was amplified by using pre-designed primers harboring the desired mutation. The oligonucleotides that were employed are listed in the **Supplementary Table 1**. DNA sequencing verified the mutations. Following confirmation, the plasmids carrying mutations were digested with XbaI (pSKPPUS-GFP) restriction enzyme from New England Biolabs to liberate the linearized plasmid. This linearized plasmid (pSKPPUS-*CaMDR1*-GFP) checked on agarose gel and then directly used to transform into *S. cerevisiæ* AD1-8u^−^ cells by employing the well-established LiAc technique (Erwin *et al*. 2007). Transformants were selected using SD-Ura^−^ agar plates (Ritu *et al*. 2007). Strains generated and used in that study are listed in the **Supplementary Table 2**.

#### Confocal microscopy

Exponential phase cells of AD1-8u-expressing GFP tagged protein variants at their C-terminal were extracted and rinsed with 1x Phosphate Saline Buffer (PBS) before being examined under Nikon A1 confocal laser microscope with a 60x oil immersion objective lens.

#### Spot dilution growth assay

For the serial dilution spot assay, after growing overnight, yeast were suspended in 0.9% saline solution at a final OD_600 nm_ of 10^6^ cells/mL at 600 nm and then diluted serially five times. A 4-μL aliquot from each dilution was put on YEPD agar plates either with or without the xenobiotic. Plates were kept at 30 °C for 48 hours (Mukhopadhyay *et al*. 2002) and final image were taken by using BIO-RAD ChemiDoc^TM^ XRS+ system.

#### MIC assay

On YEPD agar plates, yeast were cultivated overnight. The cells were then resuspended in YEPD medium to achieve a final OD_600 nm_ of 0.001 or 10^4^ cells/mL. In 96-well plates, drug dilutions were made using two-fold serial dilutions (100 μL of YEPD + 100 μL of drug with medium) up to 10^th^ well to keep one column for drug control and 100 μL cells of 0.001 OD were inoculated in each well, with last column left as control. Plates were incubated for 48 hours at 30°C and readings were taken by MIC plate reader at absorbance 595 nm. Then MIC_80_ value were determined for each strain that indicate the 80% drop of cell density relative to its drug-free control and heat map is generated using 2-color code in MS-Excel.

#### Nile red accumulation assay

Cells were grown till exponential phase from overnight culture in YEPD broth. Then 0.25 OD_600 nm_ cells was harvested and washed with PBS and resuspended in dilution medium (containing 1/3 YEPD and 2/3 water) as described (Ivnitski-Steele *et al*. 2009). Then cells were incubated at 30 °C with 200 rpm for 30 min after adding Nile red at a final concentration of 7 μM. Afterwards, cells were collected using Eppendorf Centrifuge and washed thrice with PBS before analysis on a BD FACSLyric^TM^ flow cytometer. Ten thousand cells were used for each strain to detect the geomean fluorescence intensity within the cells. The data was analyzed with inbuild BD FACSuite software. Finally, the histogram was plotted using fluorescence intensity value in percentage versus host cell control (AD1-8u^−^) and wild type CaMDR1-GFP and mutant variants of CaMDR1-GFP using the Graphpad prism software.

#### WebLogo generation

Sequences were downloaded from PFAM web-server using 192 “seed” sequences from the MFS_DHA1 subfamily (PF07690). Sequence alignment was done using the Jalview 2.11.1.4. Weblogo was generated by using web-based application WebLogo 3 using Chemistry color coding (polar-green; neutral-purple; basic-blue; acidic-red; hydrophobic-blue). At the bottom of Y-axis, numbers denote amino acid residue number of *CaMDR1* (123-512).

#### Generation 3D homology model of *Ca*Mdr1

3D model of the outward-facing conformation of *Ca*Mdr1, covering residues 110 to 544, was built on YajR, an *E. coli* proton-driven MFS antiporter crystallized in this conformation (Jiang *et al*. 2013) PDB code 3WDO. Primary sequences alignment of CaMdr1 and YajR (UniprotKB Q9URI1 and P7726, respectively) was done using AlignMe (Stamm *et al*. 2014). The alignment was used to manually superimpose the IF model of *Ca*Mdr1 with the crystal structure of YajR with Pymol (Version 2.5.0 Schrödinger, LLC.) and then submitted to Modeller (Webb and Sali 2021) that generated 20 models among which the more representative was selected manually. cytoplasmic and extracellular membrane limits as defined by the PPM server (https://opm.phar.umich.edu/ppm_server).

#### Statistical analyses

All plots were made using either GraphPad Prism (San Diego, CA) or MS-Excel. All data are represented as mean ± SD. Statistical analyses were performed using Student’s T-test. Differences were considered statistically significant when *p* < 0.05 (* signifies *p* value **≤** 0.05, ** signifies *p* value ≤ 0.01 and *** signifies *p* value ≤ 0.001).

## ABBREVIATIONS

MFS: Major facilitator superfamily
*Ca*Mdr1: Multidrug resistance 1 protein
DHA-1: Drug:H^+^ antiporter-1 subfamily
TMD: Transmembrane domain
TMS: Transmembrane segment
TMH: Transmembrane helix
ECL: Extracellular loop
ICL: Intracellular loop
CCL: Central cytoplasmic loop
CHX: Cycloheximide
4-NQO: 4-Nitroquinoline
FLC: Fluconazole
ANI: Anisomycin
CER: Cerulenin
NR: Nile red
DMSO: Dimethyl sulfoxide
WT: Wild type
PM: Plasma membrane
ABC: ATP binding cassette
SDM: Site directed mutagenesis

## ACKNOWLEDGEMENTS

We acknowledge Central Instrumental Research Facility (CIRF), Amity University Haryana, India for providing instrumental support. The collaboration with Pierre Falson, in the CNRS Research Unit « Molecular Microbiology & Structural Biochemistry, at Lyon in France, and support of SS three months stay with fellowship in Falson’s laboratory is greatly appreciated. The Senior Research Fellowship award to SS from ICMR is gratefully acknowledged. AB and RP acknowledge funding support from the Department of Biotechnology, Government of India (Grant no. BT/PR32349/MED/29/1456/2019 and BT/PR38505/MED/29/1513/2020). This work was also supported by the Centre National de la Recherche Scientifique (CNRS), Lyon University, the French Research Agency (ANR) ANR-CLAMP2-18-CE11-0002-01 to PF.

