## Supplementary material for "Spontaneous suppressors against debilitating transmembrane mutants of *Ca*Mdr1 disclose novel interdomain communication via Signature motifs of the Major Facilitator Superfamily"

Supplementary Tables 1 and 2 ; Supplementary Figures 1-9

| Primer Name | Primer Sequence |
|---|---|
| CaMdr1/F | GCATTACAGATTTTTGAGAGATAGTTTTGTTGG |
| CaMdr1/R | GCATACTTAGATCTTGCTCTCAATTTTGGTCC |
| CaMdr1 T127A/F | CAAATTTCATTTTTGGCAACTTCAGTTTATATGG |
| CaMdr1 T127A/R | CCATATAAACTGAAGTTGCCAAAAATGAAATTTG |
| CaMdr1 G140D/F | GCAGTTTATACCCCTGATATTGAAGAATTAATGC |
| CaMdr1 G140D/R | GCATTAATTCTTCAATATCAGGGGTATAAACTGC |
| CaMdr1 R184A/F | GCTATATTTGGTGCTACATCCATATATATC |
| CaMdr1 R184A/R | GATATATATGGATGTAGCACCAAATATAGC |
| CaMdr1 D235H/F | GCAAGTGTTGCTCATGTGGTTAAATTTTGG |
| CaMdr1 D235H/R | CCAAAATTTAACCACATGAGCAACACTTGC |
| CaMdr1 F216A/F | GTATATTGAGAGCCTTGGGTGGATTC |
| CaMdr1 F216A/R | GAATCCACCCAAGGCTCTCAATATAC |
| CaMdr1 G230A/F | GGCTACTGGTGCTGCAAGTGTTGCTGATGTGG |
| CaMdr1 G230A/R | CCACATCAGCAACACTTGCAGCACCAGTAGCC |
| CaMdr1 G260A/F | GTGGTCCTAGTTTTGCTCCATTCTTTGGTTC |
| CaMdr1 G260A/R | GAACCAAAGAATGGAGCAAAACTAGGACCAC |
| CaMdr1 Y378F/F | CGAAGTTTTCCCAATTTTCTTCGTTGGAGTTAAAC |
| CaMdr1 Y378F/R | GTTTAACTCCAACGAAGAAAATTGGGAAAACTTCG |
| CaMdr1 Y378A/F | CGAAGTTTTCCCAATTGCTTTCGTTGGAGTTAAAC |
| CaMdr1 Y378A/R | GTTTAACTCCAACGAAAGCAATTGGGAAAACTTCG |
| CaMdr1 Y378T/F | CGAAGTTTTCCCAATTACTTTCGTTGGAGTTAAAC |
| CaMdr1 Y378T/R | GTTTAACTCCAACGAAAGTAATTGGGAAAACTTCG |
| CaMdr1 A435T/F | GTGTTTATTCCAATTACCATTGTTGGTGGTATC |
| CaMdr1 A435T/R | GATACCACCAACAATGGTAATTGGAATAAACAC |
| CaMdr1 L480A/F | GATTTTCCAAACAGCATTCAATTTCATGGG |
| CaMdr1 L480A/R | CCCATGAAATTGAATGCTGTTTGGAAAATC |
| CaMdr1 F497A/F | TATATTGCTTCA GTTGCTGCATCAAATGATTTG |
| CaMdr1 F497A/R | CAAATCATTTGATGCAGCAACTGAAGCAATATA |
| CaMdr1 P528H/F | GGCTACCCCTGAATATCATGTTGCTTGGGGTAG |
| CaMdr1 P528H/R | CTACCCCAAGCAACATGATATTCAGGGGTAGCC |
| CaMdr1 P528A/F | GGCTACCCCTGAATATGCAGTTGCTTGGGG |
| CaMdr1 P528A/R | CCCCAAGCAACTGCATATTCAGGGGTAGCC |

**Supplementary Table 1. Oligonucleotides used in the study**.

| Strains | Genotype or description | Source |
|---|---|---|
| AD1-8Ura- | (Mata,pdr1-3,ura3 his1, Δyor1::hisG, Δsnq2::hisG, Δpdr5::hisG,Δpdr10::hisG, Δpdr11::hisG, Δycf1::hisG, Δpdr3::hisG, Δpdr15::hisG) | Kenjirou *et al*. 2001 |
| AD-MDR1-GFP | AD1-8u- cells harboring MDR1-GFP ORF integrated at PDR5 locus | Ritu *et al*. 2007 |
| **AD-MDR1 cells carrying the following mutation in MDR1-GFP ORF and integrated at PDR5 locus** (Redhu *et al*. 2018). | | |
| TMH1: I123A, T127A, T128A, S129A, Y131A, M132A, D147A | | |
| TMH2: L161A, F162A, V163A, Y166A, G167A, R184A | | |
| TMH3: Y188A, T191A, 195A, Q199A | | |
| TMH4: R215A, F216A, F220A, S223A, P224A, T228A, G229A, G230A | | |
| TMH5: W249A; P257A | | |
| TMH6: I283A, T294A, L295A | | |
| TMH7: V353A, Y360A, I361A, V364A, Y365A, L368A, Y369A, L370A, F371A, F372A | | |
| TMH8: Y394A, V398A, I399A, F406A, Y408A, P410A, E429A | | |
| TMH9: G438A, G439A, I448A, | | |
| TMHS10: A466G, 470G, F474A, I476A, F477A, Q478A, L480A | | |
| TMH11: V495A, N500A, R504A, S509A | | |
| **AD-MDR1 cells carrying the following mutations in MDR1-GFP ORF and integrated at PDR5 locus. This study** | | |
| T127A, G140D, T127A-G140D, R184A, D235H, R184A-D235H, F216A, G260A, F216A-G260A, A435T, L480A, L480A-A435T, G230A, P528H, G230A-P528H, Y378A, Y378T, Y378F, G230A-Y378A, G230A-Y378T, Y378A-P528H, P528A, F497A, G230A-F497A | | |

**Supplementary Table 2. Yeast strains used and generated in this study**.

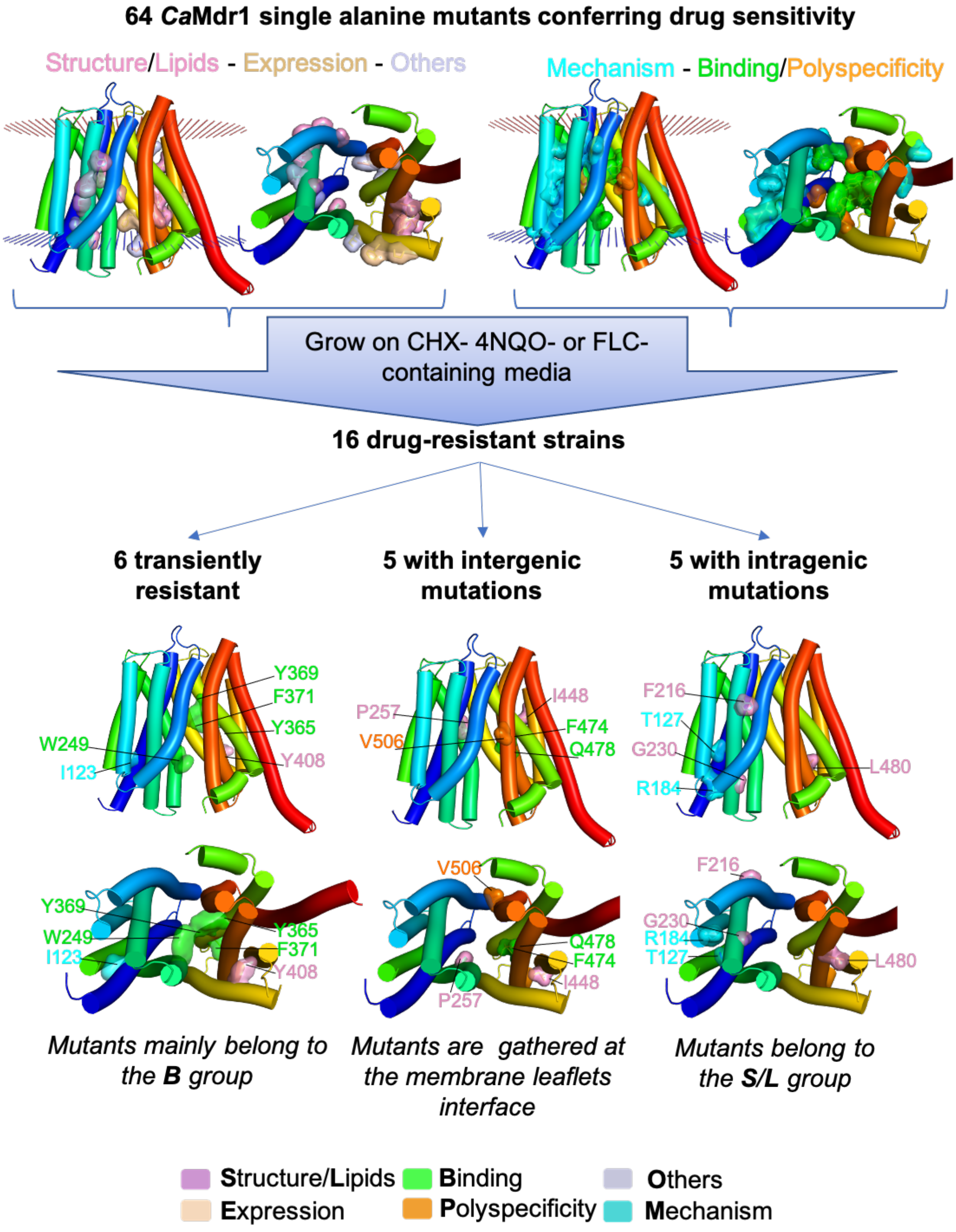

**Supplementary Figure 1. Strategy of drug-resistant strain isolation and mapping of residues screened**. GlpT-based 3D model of *Ca*Mdr1 in inward-facing conformation (Redhu *et al*. 2016) optimized with Modeller in this study. TMHs limits are defined with the OPM server (https://opm.phar.umich.edu/ppm_server). The 3D model is shown in filled cylinders colored in rainbow from the N-terminus (blue) to the C-terminus (red). Screened residues are shown in surface and colored in respect of their group as indicated and previously defined (Redhu *et al*. 2018).

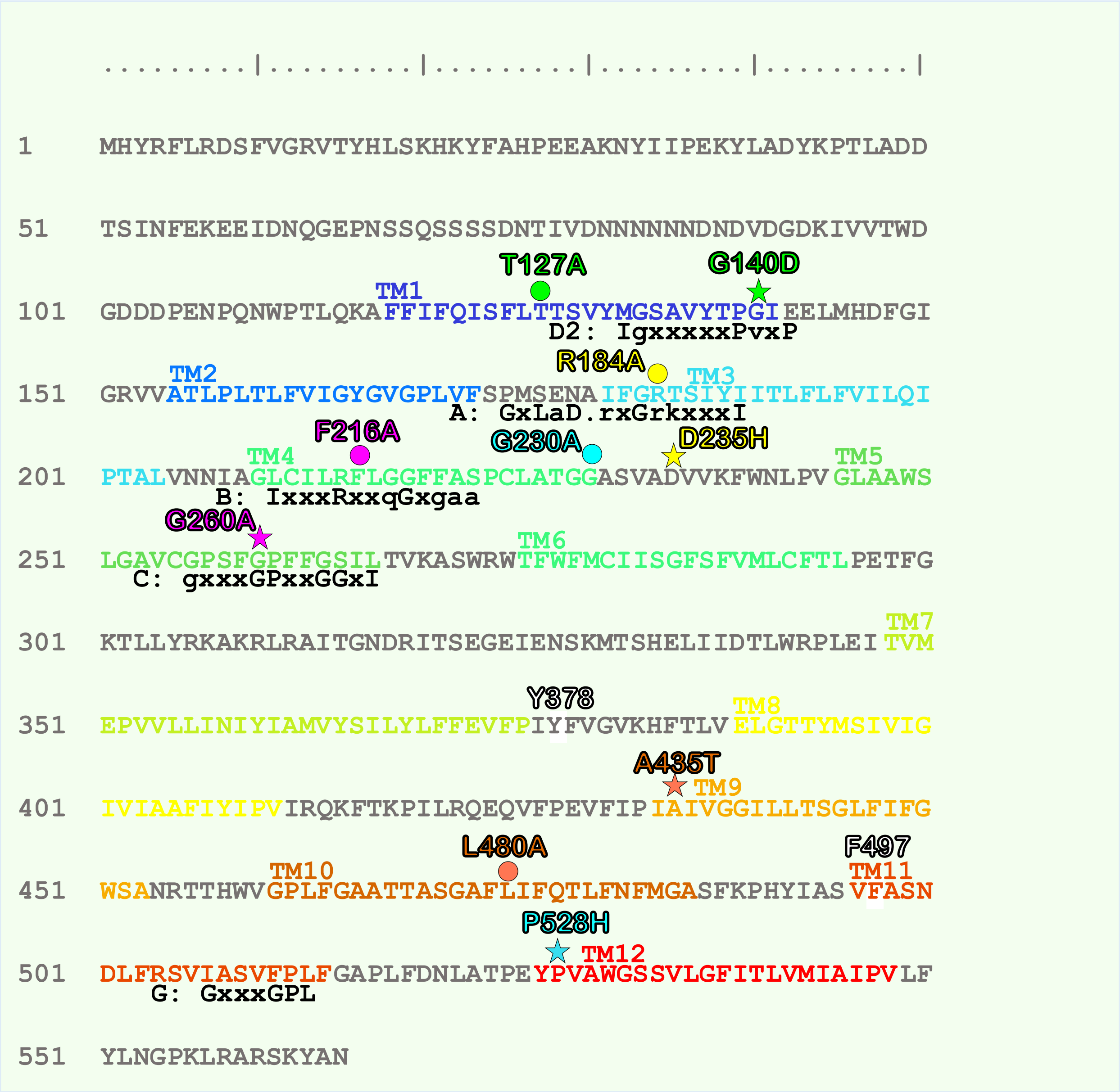

**Supplementary Figure 2.** ***Candida albicans*** **Multidrug resistance protein 1 (Uniprot # Q9URI1) primary sequence and localization of conserved motifs and mutants of the study**. TMHs are rainbow-colored in respect of the GlpT-based 3D model (Redhu *et al*. 2016) optimized here. TMHs limits are defined with the OPM server (https://opm.phar.umich.edu/ppm_server). Each couple of primary-debilitating (circle) and secondary-rescuing (star) transport mutants are colored as in Figure 1. Signature motifs of proton-dependent multidrug efflux systems are defined as in Paulsen *et al*. 1996 (see Supplementary Figure 3).

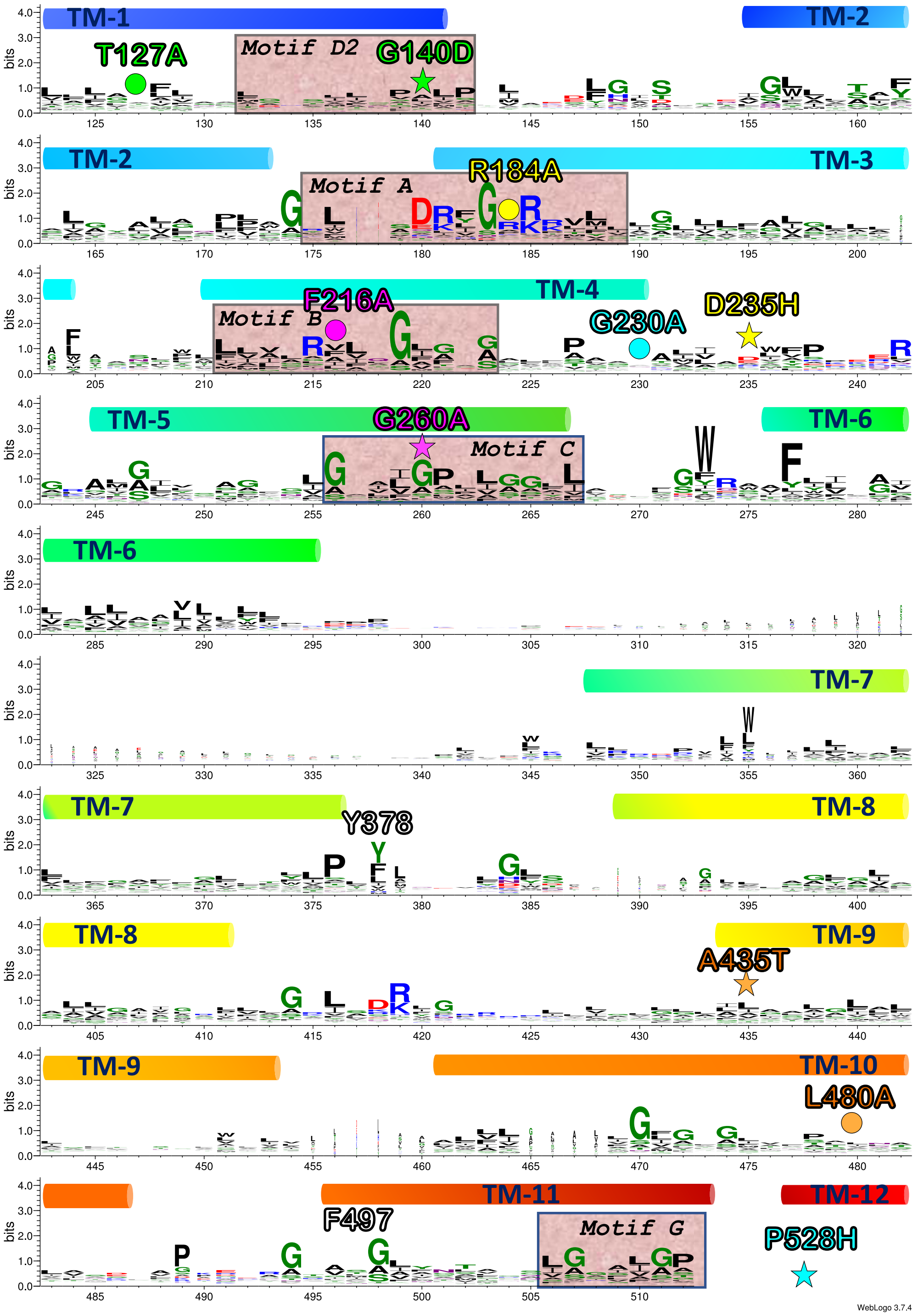

**Supplementary Figure 3. Weblogo representation of DHA1 subfamily of MFS transporters**. Sequences are from the PFAM web server using 192 "seed" sequences from the MFS_DHA1 subfamily (PF07690). Sequence alignment is done with Jalview (2.11.1.4). Weblogo is generated by WebLogo 3. Residues are colored using the Chemistry color code: polar-green; neutral-purple; basic-blue; acidic-red; hydrophobic-black. X-axis denotes amino acid residue number of *Ca*Mdr1.

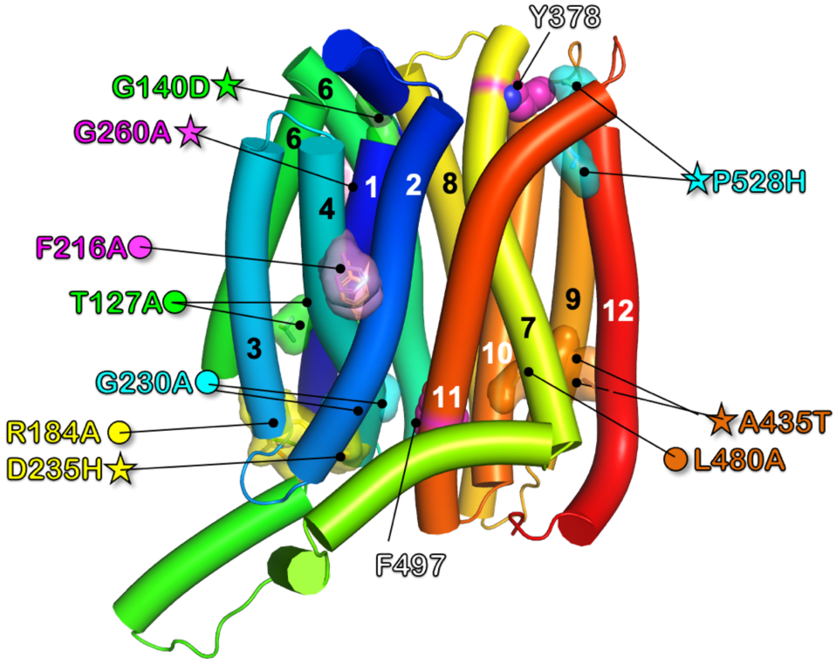

**Supplementary Figure 4. Location of critical residues in the AlphaFold 3D model of *Ca*Mdr1.** 3D model of *Ca*Mdr1 from the AlphaFold database (AF-Q5ABU7-F1-model_v1) using the UniProt entry number Q5ABU7. The picture only shows the TMH region of the model in which pairs of debilitating and rescuing mutations are displayed as in Fig. 2. Residues Y378 and F497 described later are also shown.

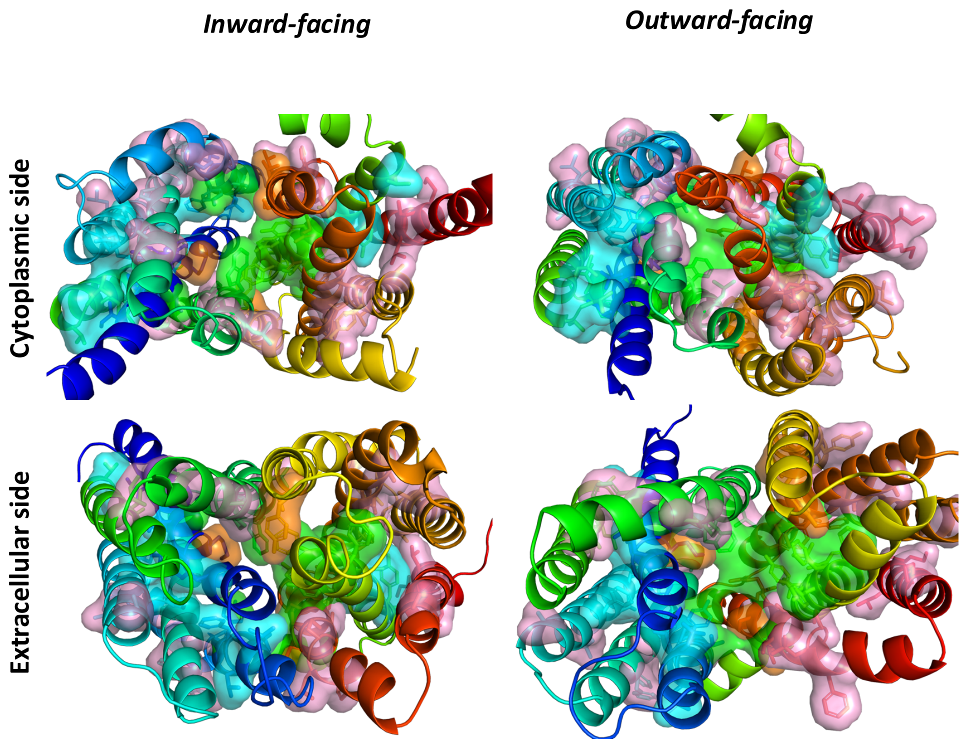

**Supplementary Figure 5. Location of critical residues in the 3D models of *Ca*Mdr1 in inward- and outward-facing conformations.** Critical residues from the alanine mutants library (Redhu *et al*. 2018) are shown in surface and stick modes and colored in respect of their impact on either the mechanism (blue), interaction with lipids and structure (pink), ligand binding (green) and polyspecificity (orange).

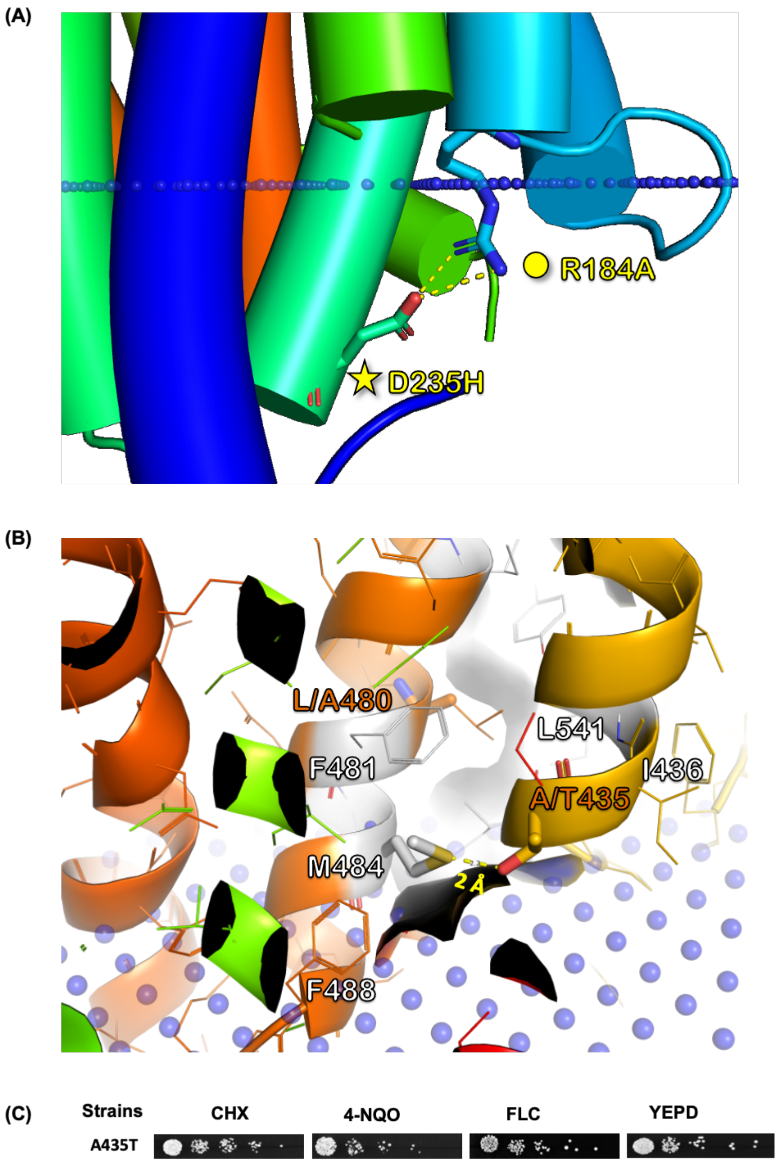

**Supplementary Figure 6. Location details of mutant and suppressor couples R184A-D235H and L480A-A435T and spot dilution assay of the A435T single mutant.** Residues are rainbow-coloured from the N- (blue) to the C-ter (red). Blue dots indicate cytoplasmic membrane limits as defined by the PPM server (https://opm.phar.umich.edu/ppm_server). (**A**) Details of the salt bridge between R184 and D235 in *Ca*Mdr1 WT (**B**) Zoom in showing the replacement of each residue and polar interaction of the OH of T435 with the sulphur atom of M484. (**C**) spot dilution assay of the A435T single mutant.

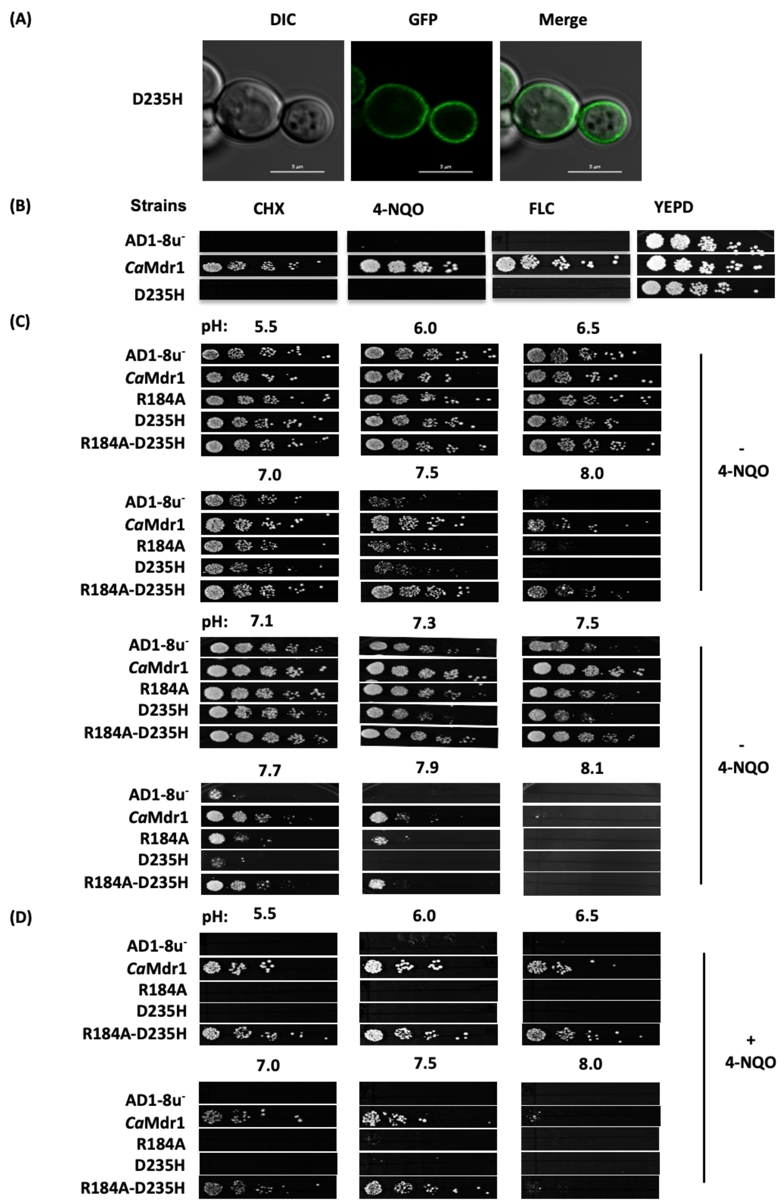

**Supplementary Figure 7. Exploration by spot dilution assays of the pH dependency of single and double mutants of the R184-D235 couple**. (**A**) Localization and (**B**) drug sensitivity of the D235H mutant. (**C**,**D**) pH dependency in respect of pH, with or without drug as indicated.

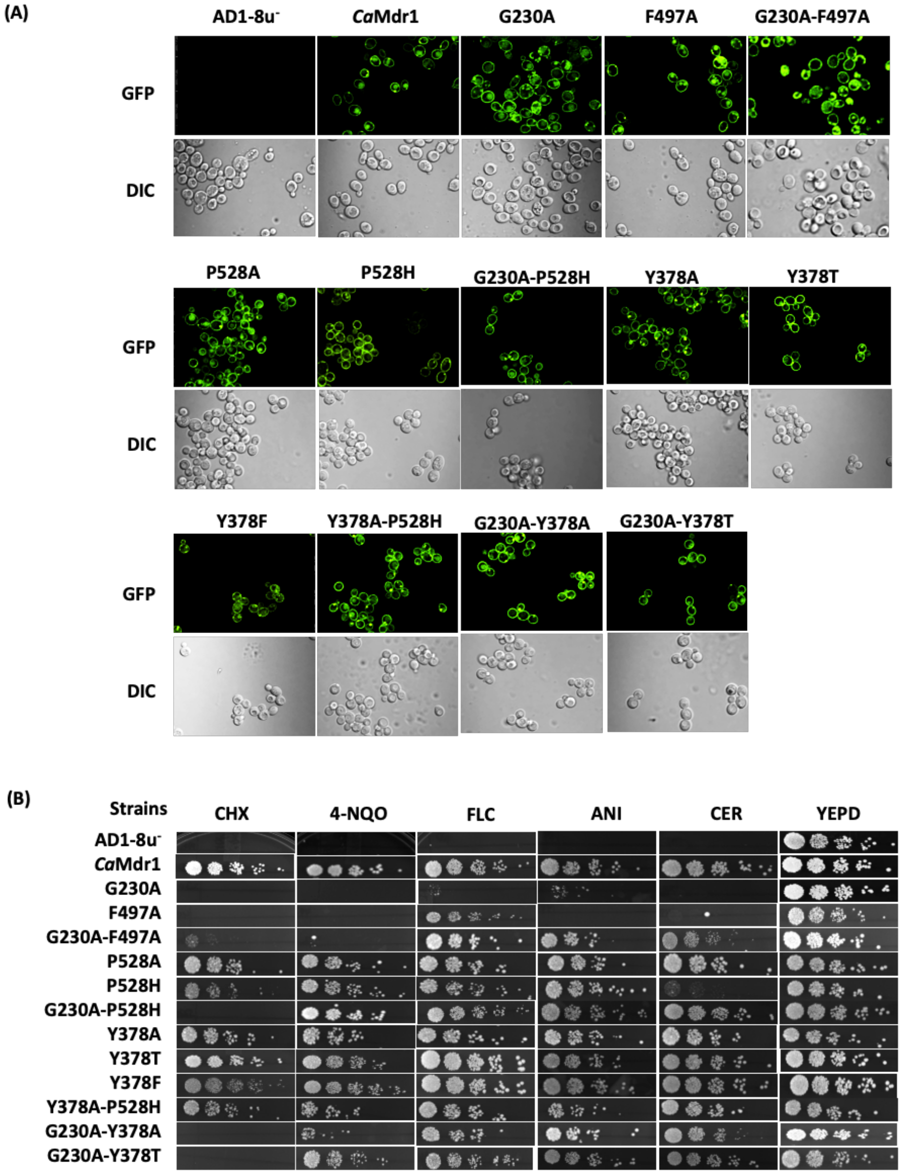

**Supplementary Figure 8. Phenotypic characterization of yeast strains expressing single secondary mutants of *Ca*Mdr1**. **(A)** Fluorescence imaging by confocal microscope showing PM localization of AD1-8u- (control), *Ca*Mdr1-GFP (WT), mutant and reconstructed suppressor with corresponding differential interference contrast (DIC) images and merged images**. (B)** Comparison of growth by spot dilution assays for mutant variants on different *Ca*Mdr1 substrates as CHX (0.1 mg/L), 4- NQO (0.15 mg/L), FLC (0.8 mg/L), ANI (10 mg/L) and CER (4 mg/L) in YEPD agar plates. Images were captured after 48 hours of incubation at 30 °C.

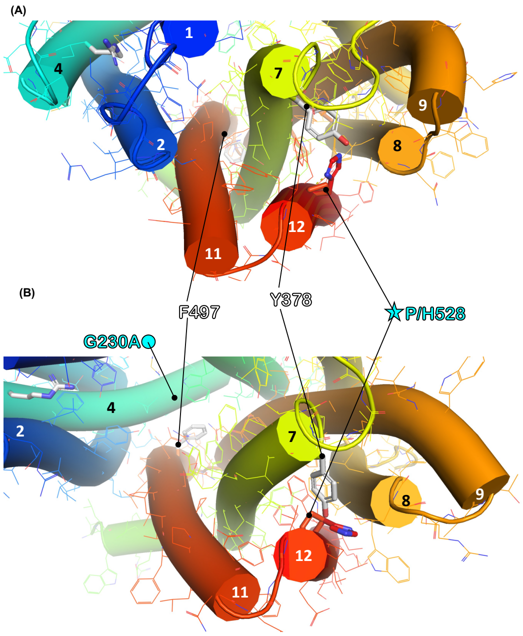

**Supplementary Figure 9. Relative position of P/H528 and Y378 in the inward- and outward-facing models of *Ca*Mdr1**. **(A)** view from the extracellular side in inward- (**A**) and outward- (**B**) facing conformations. Structural settings are as in Fig. 2.
